# Modelling assessment of the different paths of between-farm transmission of avian influenza HPAI in Northern Italy in 2021–2022

**DOI:** 10.64898/2026.09.19.752832

**Authors:** Mattia Sensi, Sara Sottile, Diletta Fornasiero, Isabella Monne, Paolo Mulatti, Andrea Pugliese

**Author notes:** These authors contributed equally to this work (Mattia Sensi, Sara Sottile).

## Abstract

We propose a stochastic epidemic model to describe the spread of avian influenza in a network of poultry farms. We consider three main paths of infection transmission between poultry farms: from nearby farms, mainly through airborne diffusion; from farms belonging to the same company, through shared veterinarians, forage providers and similar; from undetected small farms or wildlife. From the parameter estimation, based on a modified Expectation-Maximization algorithm, we infer that approximately 63% of the farms were infected from nearby ones, with an infection force declining with distance; of these, more than one third belonged to the same company of the estimated infector. About 20% were infected from premises belonging to the same company but farther away than the distance threshold of 2 km; the remaining ones from wildlife or unidentified sources. These estimates have been validated by a comparison with genetic data, available for a subset of the farms: the genetic distance between two farms identified, with high probability, as an infector-infectee pair is much lower than between random pairs. These results may help in the implementation of tailored prevention and control measures for future outbreaks.

## 1 Introduction

Avian influenza remains a major threat to animal health, food security and human health due to its capacity for cross-species transmission [1]and rapid evolutionary change [2]. Since the 2021–2022 epidemic season, Europe has experienced recurrent highly pathogenic avian influenza (HPAI) epidemic waves involving wild birds, poultry, and captive birds [3, 4]. Wild waterbirds constitute a major natural reservoir for avian influenza viruses and can contribute to their long-distance dissemination. Since control of HPAI in the wildlife reservoir is not a realistically achievable objective, control measures in domestic poultry rely on preventing introduction, ensuring early detection, and rapidly limiting secondary spread once outbreaks occur [5, 6]. This context highlights the need for quantitative modelling frameworks able to reconstruct transmission pathways, assess the relative contribution of alternative spread mechanisms, and integrate heterogeneous epidemiological, spatial, production-network, and genomic information [7].

The 2021–2022 H5N1 HPAI epidemic in Italy provides a relevant case study for farm-level epidemic reconstruction. The epidemic was largely concentrated in the densely populated poultry areas of Northern Italy, where high farm density, heterogeneous production structures, repeated opportunities for direct or indirect connections between premises, and the considerable presence of wild aquatic birds can hamper the identification of transmission pathways. During the epidemic wave epidemiological and genomic analyses suggested that multiple viral introductions were followed by extensive lateral spread in the domestic sector, highlighting the need for quantitative approaches that can jointly account for spatial proximity, temporal compatibility, and production-chain structure [8].

Over the last 20 years, avian influenza transmission has been studied using a wide variety of mathematical approaches. Spatial spread, farm density, and local clustering have been investigated using spatially explicit models and risk maps [9, 10, 11, 12], while culling strategies and their epidemiological consequences have been explored in dedicated modelling frameworks [9, 13]. Stochastic compartmental models and alternative spatial transmission kernels have also been used to describe farm-level spread in poultry populations [14, 15]. Other models have considered environmental transmission via infected faeces, viral particles, or the cocirculation of low pathogenic (LPAI) and HPAI viruses [16, 17, 18, 19]. Realistic parameter ranges are crucial for meaningful model simulations, and have been summarised in multiple reviews of HPAI transmission and dynamics [7, 20]. Although human infection with avian influenza remains uncommon, several studies have considered zoonotic spillover and transmission dynamics in humans [21, 22, 23]. Finally, the repeated introduction of avian influenza viruses into poultry farm is often linked to seasonal migration movements of wild birds. Such periodicity can be represented explicitly in mathematical modelling [12, 24, 25].

Despite these developments, reconstructing farm-level transmission during real poultry epidemics remains challenging, particularly when infection times are unobserved, several candidate infectors are simultaneously plausible, and transmission may arise from a combination of local spatial spread, production-chain connections, and external sources such as wildlife or unobserved infected premises. In this setting, genomic information can provide an independent line of evidence, but it is not always directly integrated into mechanistic reconstruction frameworks.

In this paper, we develop a stochastic model to reconstruct the spreading of H5N1 HPAI in a network of poultry farms. Using data from the 2021–2022 H5N1 HPAI epidemic in Northern Italy, we estimate unobserved infection times and probabilistic transmission pathways for outbreaks from outbreak data in which detection and removal times are observed, but infection dates and transmission sources are not. We then compare the inferred transmission structure with available genomic data from the main cluster of infected farms, using genetic distances as an independent source of evidence for the plausibility of reconstructed links.

The remainder of this manuscript is structured as follows. In Section 2, we describe the data used, the compartmental model used in our work, and the statistical methods implemented. We present our results in Section 3, and we discuss them, as well as possible research outlook, in Section 4.

## 2 Material and Methods

### 2.1 Data Availability

The data used in this study consist of multiple datasets describing poultry farms, their geographical location, epidemiological status, and genetic information. The full dataset includes 1653 commercial poultry farms farms, of which 297 were infected during the 2021–2022 winter epidemic in Northern Italy.

The available data include information on all the farms included in the study, such as unique identifiers and geographical coordinates (latitude and longitude), as well as company-level information linking farms belonging to the same production chain, including additional attributes such as production type and species. In addition, temporal epidemiological data are available, namely detection and removal dates for infected farms.

We also considered additional data related to the largest outbreak cluster, consisting of 214 infected farms. For this subset, pairwise genetic distances based on nucleotide differences between viral sequences are available [8], allowing comparison between inferred transmission links and genetic similarity.

All datasets were preprocessed to ensure consistency in identifiers and temporal formats, and were merged to construct the final dataset used for model estimation and visualization.

All analyses and visualizations were implemented in Python using standard scientific and geospatial libraries, including NumPy, SciPy, pandas, GeoPandas, and Matplotlib. The code used in this study is available at https://github.com/SaraSottile/AvianFluNetwork.git, together with synthetic data which can be used to replicate our results. Due to data privacy and confidentiality constraints, the raw data cannot be publicly shared. However, aggregated data and derived quantities supporting the findings of this study are available upon reasonable request.

### 2.2 Model Overview

We consider a fixed population of *N* farms, each of which can be in one of five epidemiological states: *S* (Susceptible), *E* (Exposed, i.e. infected but not yet infectious), *U* (infected and infectious but Undetected), *D* (infected and Detected), and *R* (Removed after culling and disinfection). The transition from *D* to *R* corresponds to the complete culling of poultry within the farm, as a consequence of the confirmed detection of avian influenza.

The model describes the infection dynamics at the level of farms, rather than individual animals, similarly, for instance, to [12, 14, 26]. A schematic representation of the compartmental structure is shown in Figure 1. Farms transition from *S* to *E* upon infection, then progress through the latent phase (*E*) to the infectious undetected state (*U*), and are eventually detected (*D*), after which control measures lead to removal (*R*) through culling of the entire poultry population within. Farms may also be repopulated and return to the susceptible state (*R → S*), or be preemptively culled (*S → R*) if they are located too close to a detected outbreak.

**Figure 1:**
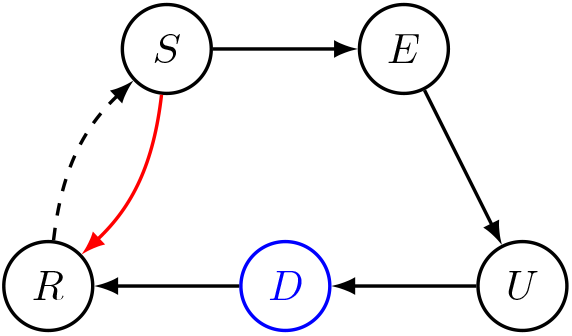
Flow diagram of the model. The probability of moving from *S* to *E* depends on the force of infection exerted on a given farm in a given day, defined in Section 2.3; the passage from *E* to *U* depends on the dormant phase of the infection; a farm moves to state *D* once the infection is spotted and confirmed, after which the poultry is culled, bringing it to state *R*. We stress that the only available data are the detection dates for each eventually infected farms (*U → D*), the dates of culling (*D → R*) and, potentially, the dates of re-population (*R → S*, as farms can be sanitised and repopulated after a culling). Additionally, farms can potentially be culled preemptively (*S → R*) if they are close to a confirmed infection.

The available observational data consist of the detection dates for each infected farm (corresponding to the transition from *U* to *D*) and the removal (culling) dates (corresponding to the transition from *D* to *R*). The dates of exposure (*S → E*) and development of infectiousness (*E → U*) cannot generally be known exactly; with our model we aim to provide reasonable estimates in particular of the date of exposure of each eventually detected farm. The time spent in the exposed state *E* has been estimated to be around 2 days [27]. This is followed by an infectious period that begins in the *U* compartment, whose duration may be highly variable with an average of 4–5 days prior to detection, and may extend for a short and variable duration into the *D* compartment until culling. Infectiousness is therefore restricted to the late pre-detection and early post-detection phases, corresponding to the *U* and *D* compartments.

In practice the transitions between *E* and *U* cannot be directly ascertained from the available epidemiological data. For this reason, the model we used does not explicitly reconstruct the onset of infectiousness as a separate observable event. Instead, infectiousness is described through the continuous infectivity profile introduced in Section 2.3, which progressively increases between the inferred exposure time and the detection date. As a consequence, the effective reconstruction problem is reduced to the estimation of the exposure time *E* given the observed detection time *D*, while the intermediate compartment *U* is implicitly represented through the infectivity dynamics.

### 2.3 Force of Infection

Each eventually infected farm *j ∈ {*1, 2, …, *N}* is associated with a detection date, denoted by *D*_*j*_.

Consistently with the compartmental durations described in Section 2.2 and with the protocols [28, 29], we define a temporal window for the infection time as

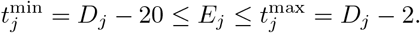

leaving a minimum of 2 days between infection and detection (corresponding to very short incubation and pre-detection periods) and 20 days to allow for very large variability in the incubation phase, detection delays, and uncertainty in the underlying processes. This interval is used as a constraint within the inference algorithm to estimate the most likely infection time and source.

#### Infectiousness profiles

We assume that, before detection, infectiousness grows exponentially, in parallel with the exponential growth in the number of infected animals in the farm. Specifically, the relative infectiousness of a farm *k* days before detection is assumed to be equal to *γe*^*−γt*^. Correspondigly, for each farm *k*, we introduce a function *ϕ*_*k*_(*t*) describing the infectiousness of the *k*-farm in day *t*, given by

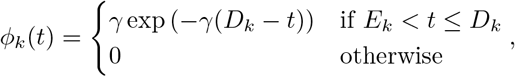

since a farm cannot be infectious before *E*_*k*_ + 1, the day after being infected,

In the estimation, we set *γ* = 1*/*10; corresponds to a characteristic timescale of approximately 10 days between infection and detection inferred from the epidemiological data. To assess the robustness of our results to this assumption, we also perform a sensitivity analysis using *γ* = 1*/*5, reported in Appendix A.

#### Force of infection

We assume that the force of infection acting on farm *j* at time *t* is the sum of three components:

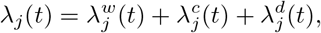

whose transmission mechanisms are explained below.

#### Wildlife and unobserved sources

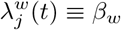

is a constant (to be estimated) accounting for infection from wild birds or unreported farms. We assume for lack of specific information, that there is a constant background infection force throughout the the whole duration of the epidemic.

#### Company network transmission

We assume that contacts between farms belonging to the same company may be a considerable vector of disease spreading, as these farms realistically share veterinarians, forage providers, etc. Precisely, we assume that the force of infection that farm *k* exerts on farm *j* through the shared production chain is

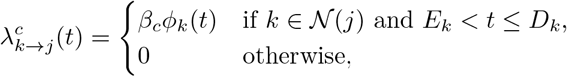

where *β*_*c*_ is a constant (to be estimated) and *N*(*j*) denotes the set of farms belonging to the same company as farm *j*.

In words, we consider that infection transmission could occur from each farm in the same network that at time *t* had already been exposed, but had not yet been detected. Its contribution is then weighted according to the infectiousness profile discussed earlier.

One can then define the overall force of infection on farm *j* through the shared network as

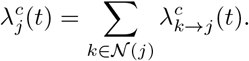

#### Distance-based transmission

The third source of transmission comes from farms located nearby, assuming that viral particles, especially present in the faeces of the animals, could get dispersed by wind or similar mechanisms. We assume that this force of infection decreases with the distance between farms, and becomes 0 beyond a given distance *d*_max_. Precisely, we use the form

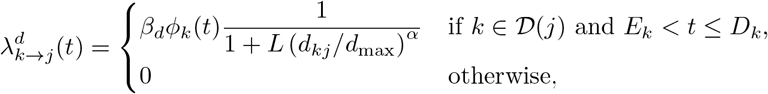

where *β*_*d*_ and *α* are constants (to be estimated), *D*(*j*) denotes the set of farms within a radius *d*_max_ from farm *j*, and *d*_*kj*_ is the distance in km between farms *k* and *j*.

The parameter *L ≫* 1 controls the decay of transmission at large distances, strongly suppressing the contribution of farms close to the cutoff distance *d*_max_. In particular, when *d*_*kj*_ = *d*_max_, the kernel contribution becomes approximately proportional to 1*/L*, although its effective magnitude still depends on the multiplicative scaling parameter *β*_*d*_. In the following, we fix *L* = 10^3^, so that the force of infection at the cut-off distance is around one thousandth of the value that holds very close to the source of infection. As for *d*_max_, we use 2 km, that is in the range used in [26]to characterise short-term transmission. In Appendix A, we consider *d*_max_ = 1.5 km to test the sensitivity of the results to this assumption.

From this expression, we then obtain

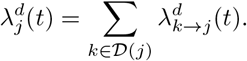

If a farm is both in *N* (*j*) and *D*(*j*), i.e. it belongs to the same company as farm *j and* it is located within *d*_max_ km of farm *j*, we allow both transmission mechanisms to contribute to the force of infection. This reflects the idea that proximity and company-related contacts represent distinct epidemiological pathways, each providing an independent contribution to the overall infection hazard.

### 2.4 Parameter estimation and simulation framework

We exploit the observed detection dates *D*_*j*_ to perform parameter estimation within a discrete-time framework with daily resolution. All farms are observed over a common time window defined from the earliest plausible infection time to the latest detection date (from late October 2021 to mid January 2022).

As an initial step, infection times are approximated as *E*_*j*_ *≡ D*_*j*_ *−* 15 for all *j* to let the algorithm explore a wide range of distances between exposure and detection. We then iteratively refine both the infection times and the model parameters.

Given the parameter set

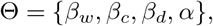

and taking the infection times *E*_*j*_ as known, we define the likelihood as

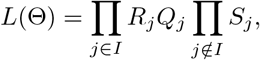

where *I* denotes the set of infected farms,

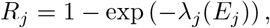

is the probability that farm *j* becomes infected at time *E*_*j*_ (conditional on not having been infected before),

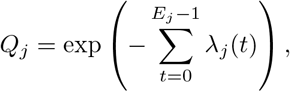

is the probability that farm *j* remains uninfected up to time *E*_*j*_ *−* 1, and

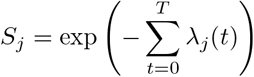

denotes the probability that a non-infected farm remains susceptible throughout the entire observation period, where *T* is the final observation time.

Parameter estimation is performed by maximizing the likelihood using a bounded L-BFGS-B optimization scheme, with parameters initialised randomly within biologically plausible ranges.

Given an estimate of Θ, we compute, for each farm *j*, the probability of infection at each admissible time 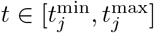, as defined in Section 2.3, according to

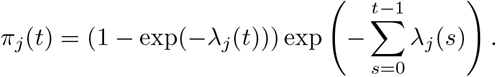

The infection date is then updated as *E*_*j*_ = arg max_*t*_ *π*_*j*_(*t*).

Using the updated infection times, parameters are re-estimated and the procedure is iterated until convergence, defined as stability of the infection dates between successive iterations, or until a maximum of 500 iterations is reached. This procedure corresponds to a classification-type expectation–maximization algorithm, where infection times are treated as latent variables.

To ensure computational efficiency, all quantities are precomputed and stored in matrix form. In particular, spatial distances and company-based connections are encoded as adjacency structures, allowing the force of infection to be evaluated efficiently through matrix operations.

To assess robustness with respect to initialization, the estimation procedure is repeated over multiple random seeds, and results are aggregated across runs.

The model is estimated multiple times using different random seeds. We define the aggregated parameters as

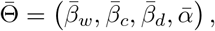

where each component is computed as the median across seeds. The corresponding estimated values are reported in Table 1 together with summary statistics, indicating the variability among estimates obtained using different seeds.

**Table 1:** Parameter estimates.

| Parameter | Median | Mean | SD |
| --- | --- | --- | --- |
| $\beta_w$ | $1.41 \times 10^{-4}$ | $1.41 \times 10^{-4}$ | $1.40 \times 10^{-7}$ |
| $\beta_c$ | $7.12 \times 10^{-4}$ | $7.11 \times 10^{-4}$ | $5.36 \times 10^{-6}$ |
| $\beta_d$ | 110.64 | 111.61 | 8.86 |
| $\alpha$ | 0.61 | 0.61 | $5.87 \times 10^{-2}$ |
| Duration ($E \rightarrow D$) | 6 | 7.83 | 5.83 |

Similarly, for each infected farm *j*, we obtain multiple estimates of the infection time;we define the consensus estimate *E*_*j*_ of the infection time as the median:

We use these estimates to obtain the duration of the time between exposure and detection

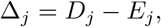

where *D*_*j*_ denotes the observed detection date.

We remark that, while only farms eventually detected as infected are retained for the construction of the transmission matrix (see below), all farms, including non-infected ones, are included during parameter estimation, as detailed above.

### 2.5 Construction of the Infection Probability Matrix

We construct a directed infection probability matrix describing the most likely transmission pathways between farms. For a given farm *j*, estimated to have been infected at time *E*_*j*_, we evaluate the force of infection at time *t* = *E*_*j*_ and decompose it into contributions from all potential sources. Let *λ*_*j*_(*E*_*j*_) denote the total force of infection acting on farm *j* at the inferred infection time, and let 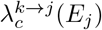 and 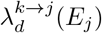 denote the contributions from farm *k* through the company and distance transmission components, respectively.

The probability that farm *k* infected farm *j* is defined as

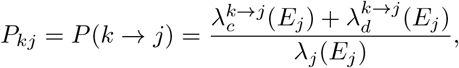

while the probability of wildlife-mediated infection is

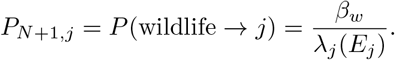

In this way we obtain a matrix *P* of size (*N* + 1) *× N*, where *N* denotes the number of infected farms included in the reconstruction. Rows correspond to potential infectors (infected farms and wildlife), while columns correspond to infected farms. Diagonal entries are set to *P*_*jj*_ = 0, since self-infection is excluded by construction. Each column *j* therefore represents a probability distribution over the possible infection sources of farm *j*:

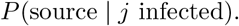

Similarly, for each farm, one can obtain the probability of having been infected from farms in the same network company or from farms nearby as

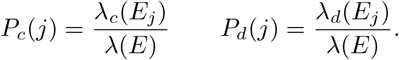

It turns out that an important source of infections are farms located nearby *and* belonging to the same company. Thus, we define *DN* (*j*) := *D*(*j*) *∩ N* (*j*) as the set of companies beloging to both *D*(*j*) and *N* (*j*), and

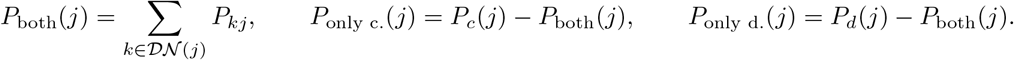

This representation allows us to extract several quantities of epidemiological interest, including the identification of the most likely infector for each farm, the quantification of wildlife-mediated infections, and the reconstruction of the inferred transmission network.

### 2.6 Visualization

The spatial visualization provides a dynamic representation of the reconstructed epidemic progression. Farms are represented as nodes positioned according to their geographical coordinates, while edges connect farms corresponding to the reconstructed transmission links. Node colors indicate the epidemiological state of each farm, and newly infected farms are highlighted to facilitate the visualization of the epidemic progression.

The visualization displays, for each day, all possible infection links identified by the transmission probability matrix, with thicker lines corresponding to higher probabilities. Link colors distinguish between company-related and distance-based transmission pathways. The visualization is generated over a daily time grid, allowing the temporal evolution of the reconstructed transmission network to be followed throughout the epidemic. Farms for which wildlife is inferred as the most likely source are not associated with a farm-to-farm transmission link.

Each time step is rendered as an individual frame, including a geographic basemap. The sequence of frames is then assembled into a video, providing a dynamic representation of the epidemic spread. The resulting visualization is available in .mp4 format in the public repository https://github.com/SaraSottile/AvianFluNetwork.git.

## 3 Results

### 3.1 Parameter estimation and epidemic reconstruction

The estimated parameter values, as well as the variability in the estimates, resulting from different starting seeds, are reported in Table 1

Appendix A.1 summarises the distribution of the estimated parameters across runs for different values of *d*_max_ and of *γ*.

A total of 297 farms were infected and eventually detected. The median inferred transition times between exposure and confirmation (*E → D*) has been of 6 days, consistently with literature estimates [27], although the starting point of the estimation procedure was set to a much larger value (15 days).

Of the 297 infected farms, we estimated the relative contribution of four possible transmission mechanisms: transmission from farms belonging to the same company, transmission from nearby farms, transmission from farms that are both nearby and belong to the same company and transmission from wildlife. For each infected farm, the dominant transmission mechanism was identified as the mechanism with the largest contribution to the estimated infection probability. The resulting classification is reported in Table 2.

**Table 2:** Distribution of the 297 infected farms according to the dominant transmission mechanism.

| Dominant mechanism | Number of farms |
| --- | --- |
| Only same company | 59 |
| Only distance | 124 |
| Distance and same company | 64 |
| Wildlife | 50 |
| Total | 297 |

Overall, distance-based transmission was the dominant mechanism for 188 farms (63%), either through distance alone (124 farms, 42%) or through farms that were both nearby and part of the same company (64 farms, 22%). Transmission attributed exclusively to farms belonging to the same company accounted for 59 farms (20%), while 50 farms (17%) were attributed to wildlife. Among the 64 farms for which both distance and same-company transmission contributed to the dominant mechanism, 63 are in the group of high probability, while 1 is in the group with medium probability. Thus, transmission from nearby farms represented the predominant mechanism, while transmission involving farms belonging to the same company also accounted for a substantial fraction of infections.

As a complementary analysis, we then considered the most likely individual infector identified for each infected farm. The inferred infector was classified according to whether it was only associated with the same company, only with the distance-based network, with both networks, or was explicitly identified as wildlife. Figure 2 reports the distribution of the corresponding best-infector probabilities within these four categories. The probability associated with the inferred best infector was high (above 50%) for almost all farms in the distance-based categories: 119 out of 124 farms (96.0%) in the only-distance category and 61 out of 64 farms (95.3%) in the distance-and-same-company category. In contrast, the wildlife category showed a more balanced distribution, with 50 farms (45.9%) having high probability and 59 farms (54.1%) having medium (between 10 and 50%) probability. No farms were classified as low probability (below 10%). No farm had a best infector belonging exclusively to the same-company network. In Appendix B, we describe the uncertainty in source attribution through different measures.

**Figure 2:**
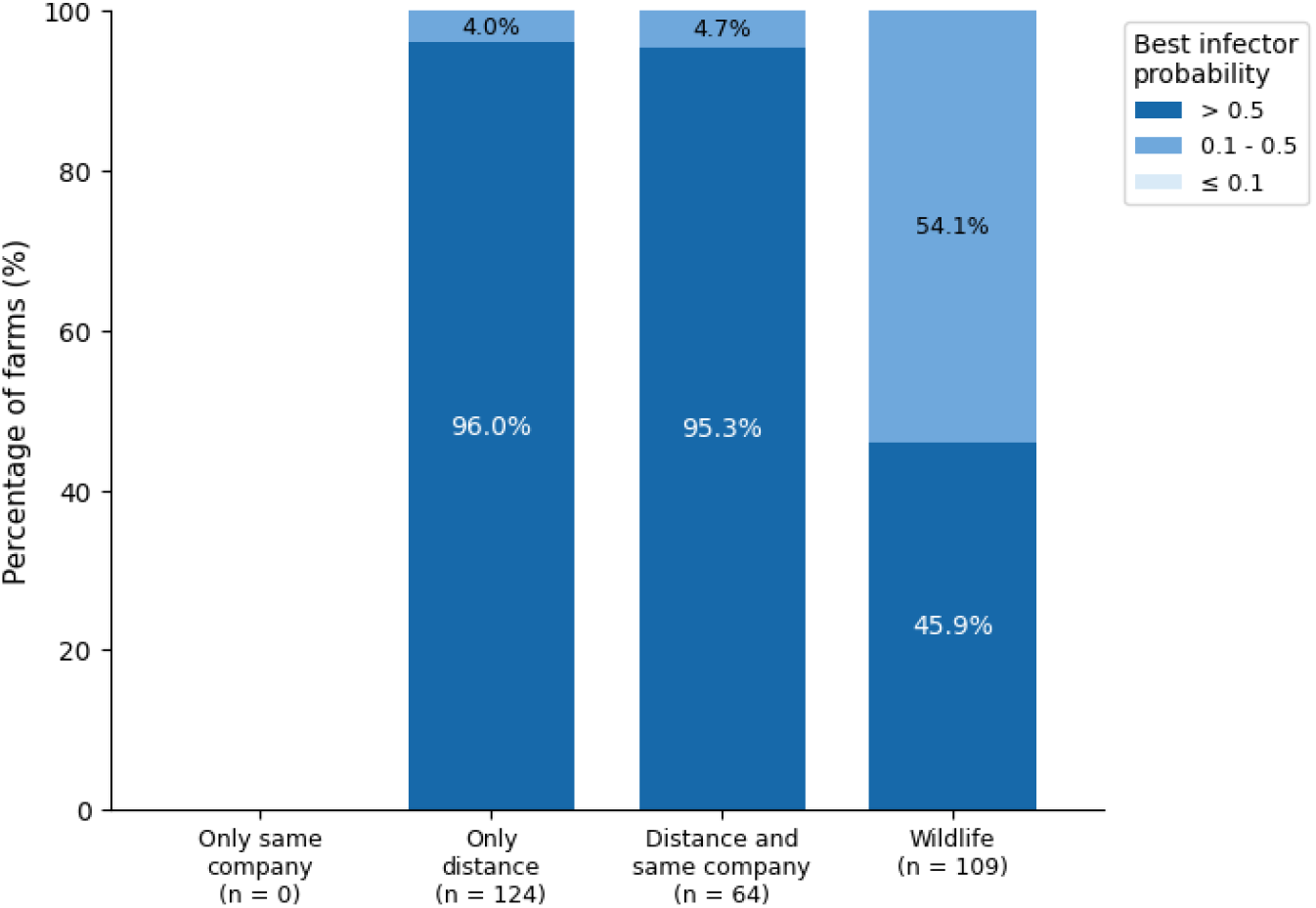
Classification of the most likely infector for the four transmission categories. Bars show the number of infected farms, subdivided according to the probability associated with the inferred best infector. High probability corresponds to values greater than 0.5, while medium probability corresponds to values between 0.1 and 0.5. No farms were classified as low probability.

The two classifications therefore provide complementary information. The dominant-mechanism classification gives a clearer and more consistent picture, with all farms falling into the high class under the 50% threshold. The best-infector analysis instead shows more variation, especially for wildlife, where a substantial proportion of cases falls into the medium probability class. This suggests that the overall transmission mechanism can be identified with high confidence, while the identification of the specific source may be less certain. In Appendix A.2,we provide a detailed description of the dominant transmission mechanism classification and of the additional classification based on the most likely infector.

### 3.2 Global comparison with genetic distances

The directed transmission matrix *P* (*k → j*), built in the previous Subsection, was then compared with the matrix of pairwise genetic distances. The comparison was restricted to the subset of farms present in both datasets. A total of 210 farms had available genetic information, resulting in 43, 890 ordered pairs (*k, j*) included in the analysis. The wildlife source was excluded, as no genetic sequence is associated with it and therefore no genetic distance can be defined.

For each ordered pair (*k, j*), the transmission probability *P* (*k → j*) was matched with the corresponding genetic distance. We first evaluated the overall association using Spearman correlation, which resulted in a value close to zero and not statistically significant (*ρ* = *−* 0.0009, *p* = 0.857). This indicates that no clear global relationship emerges when considering all possible pairs, likely due to the highly unbalanced nature of the data, where most pairs are associated with very low transmission probabilities.

Using the same probability classes introduced above, we performed the analysis on all inferred transmission pairs, rather than only the dominant source associated with each infected farm. Transmission pairs were therefore stratified into high (*P* (*k → j*) *>* 0.5), medium (0.1 *< P* (*k → j*) *≤* 0.5), and low (*P* (*k → j*) *≤* 0.1) probability classes. For the purpose of this pairwise analysis, the wildlife source was excluded because no farm-to-farm transmission pair can be defined for this source and no corresponding genetic distance is available. All remaining farm-to-farm pairs were classified according to their individual transmission probabilities. The resulting distributions of genetic distances are summarised in Table 3.

**Table 3:** Genetic distances stratified by inferred transmission probability.

|  | High probability | Medium probability | Low probability | Most likely infector |
| --- | --- | --- | --- | --- |
| $n$ | 126 | 109 | 43,655 | 132 |
| Mean distance | 7.92 | 13.21 | 21.43 | 8.08 |
| Median distance | 5 | 11 | 21 | 5 |

It can be seen from Table 3 that transmission pairs assigned to the high-probability class exhibited the smallest genetic distances, with median value of 5. Medium-probability pairs show intermediate genetic distances (median 11), whereas low-probability pairs are associated with substantially larger genetic divergence (median 21). The same trend can be observed when considering only the most likely infector identified for each infected farm. After excluding infections attributed to wildlife, the inferred infectors show median values of 5. The sensitivity analysis is showed in Appendix A.3.

The distributions of genetic distances across the different probability classes are shown in Figure 3. The relationship between probability classes anf genetic distance is confirmed by statistical tests: both the Mann–Whitney and Kruskal–Wallis tests showed highly significant differences among probability classes (Table 4), demonstrating that transmission pairs assigned higher probabilities are characterised by significantly smaller genetic distances.

**Table 4:** Statistical comparison of genetic distances across transmission probability classes.

| Test | p-value |
| --- | --- |
| Mann–Whitney | $6.63 \times 10^{-43}$ |
| Kruskal–Wallis | $2.86 \times 10^{-56}$ |

**Figure 3:**
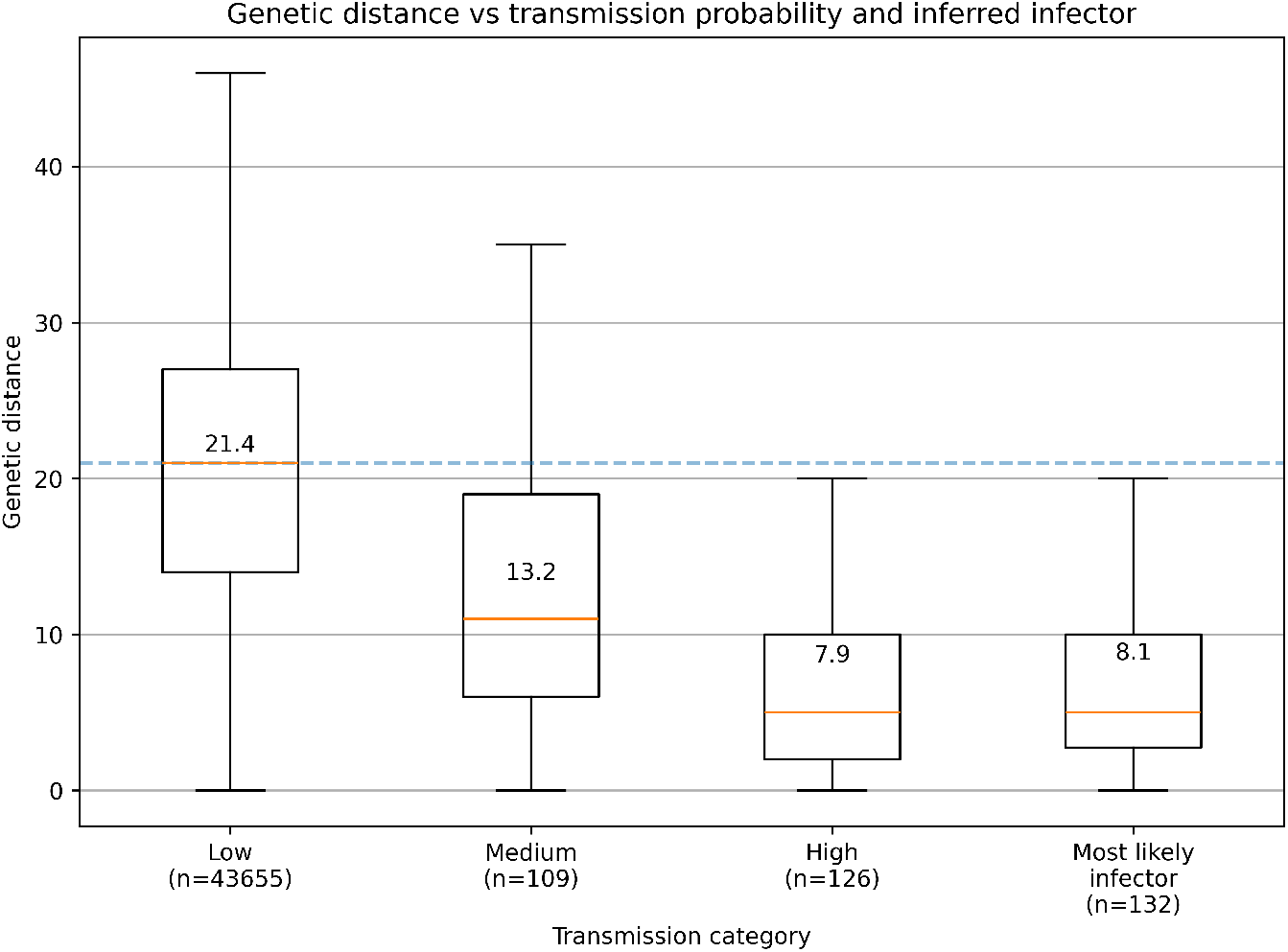
Comparison between inferred transmission probability and genetic distance. Pairs were stratified into three classes based on transmission probability: low (*P ≤* 0.1), medium (0.1 *< P ≤* 0.5), and high (*P >* 0.5). The distributions corresponding to the most likely infectors (excluding wildlife-attributed cases) are also shown. The numbers in each box represent the mean genetic distance.

Overall, these results provide independent support for the biological plausibility of the inferred transmission network. While no meaningful global correlation is observed across all possible farm pairs, the model consistently assigns the highest transmission probabilities to genetically more similar farms.

### 3.3 Spatial visualization of the epidemic dynamics

To provide a dynamic and intuitive representation of the epidemic spread, the model outputs were projected onto the geographical map of the farms. Initially, all farms are displayed as susceptible nodes. As the epidemic unfolds, nodes change color according to their epidemiological status and timing, specifically reflecting the inferred dates of exposure, the observed dates of detection, and the observed removal (culling) dates.

Susceptible farms are represented in blue, exposed farms in red, detected farms in green, and removed farms in black. Newly exposed farms are additionally highlighted using larger node sizes in order to emphasise the temporal progression of the epidemic.

This visualization emphasises both the temporal and spatial structure of the transmission process, making it possible to identify localised clusters of infection as well as the relative contribution of different transmission mechanisms. In particular, one sees the progressive emergence of geographically connected clusters interspersed with few long jumps; this visually highlights the dominant role of local spatial transmission observed in the statistical analysis.

## 4 Discussion

### 4.1 Interpretation of results

The results of this study provide a coherent picture of the transmission dynamics at the farm level, highlighting the dominant role of spatial proximity in driving the spread of infection. Differently from other analyses of highly pathogenic avian flu [12, 14, 26], we consider a special component of the transmission between farms belonging to the same company, due to likely sharing of veterinarians, suppliers, and others.

The results suggest that the two transmission pathways may act synergistically: distance-based transmission was the dominant mechanism for 188 of the 297 infected farms, including 64 farms for which both the distance-based network and the same-company network contributed to the dominant mechanism. Transmission attributed exclusively to the same-company network accounted for 59 farms.

This suggests that local transmission processes, potentially including environmental exposure, short-range indirect contacts, in particular between poultry farms of the same company, represented the dominant reconstructed component of spread.

The wildlife component accounted for a smaller proportion of infections than the distance-based mechanism, although it remained a non-negligible source of infection; however, it is unlikely to represent the dominant driver of the epidemic under the conditions considered. Still, several events of long-range transmission have been attributed by the model to this component, clearly including the earliest inferred infected farm. Although the name suggests direct transmission from wildlife, we cannot exclude transmission from unobserved sources (especially family-run farms), or unaccounted long-range transmission.

The additional analysis of the most likely infector provides further insight into the uncertainty of source attribution. When the most likely infector was associated with distance-based transmission, it was assigned to the high-probability class in more than 94% of cases. In contrast, the attribution to wildlife was more uncertain, with a substantial proportion of cases falling into the Medium class and, for *γ* = 0.2 and *d*_max_ = 2 km, some cases also falling into the Low class. This indicates that the dominant transmission mechanism can be identified more consistently than the specific source of infection.

The comparison with genetic data further supports the biological plausibility of the inferred transmission network. Although no global relationship was observed between transmission probability and genetic distance, a clear association emerged after stratifying transmission pairs according to their inferred probability. High-probability transmission pairs consistently exhibited substantially smaller genetic distances than low-probability pairs, supporting the hypothesis that the inferred strongest epidemiological links correspond to recent transmission events. This comparison was performed on the subset of data on which genetic data were available. The result supports the plausibility of an inference based on epidemiological data only.

There were two important assumptions in the analysis: first, we used a cut-off distance of 2 km for distance-dependent transmission; second, we assumed that infectiousness of a farm increases when getting closer to detection date, since the number of infected birds within a farm is expected to grow exponentially in the first phase of an outbreak; correspondingly, we assumed that infectiousness grows exponential at rate *γ* = 0.1 (day)^*−*1^, i.e. with a doubling time of around 7 days. Since both assumptions are questionable, we performed a sensitivity analysis using *d*_max_ = 1.5 km, and *γ* = 0.2 (day)^*−*1^.

Results from the sensitivity analysis (Appendix A) show that most results are robust to the choice of the parameters *d*_max_ and *γ*. Parameter estimates are similar, distance-based transmission remains the dominant mechanism for both values of *γ*, and the relationship between transmission probability and genetic distance remained remarkably consistent.

When the cut-off distance *d*_max_ is chosen smaller, the main difference is that a number of infections that were attributed to nearby farms when *d*_max_ = 2 km become attributed to wildlife (or unidentified) sources. Keeping *γ* = 0.1, the number of farms for which distance-based transmission was dominant decreased from 188 (when *d*_max_ = 2 km) to 146 (when *d*_max_ = 1.5 km), while the number attributed to wildlife increased from 50 to 83. A similar pattern was observed for *γ* = 0.2, with distance-based transmission decreasing from 184 to 142 farms and wildlife increasing from 43 to 78.

Another difference between the cases of *d*_max_ = 2 km or *d*_max_ = 1.5 km is that the median gap between the first and second most likely infectors decreased from 0.411 (*d*_max_ = 2 km) to 0.223 (*d*_max_ = 1.5 km), while the proportion of farms with *P*_1_ *>* 0.5 (i.e. the infector was identified with a probability higher than 50%) decreased from 60.6% to 47.1%. These findings indicate that reducing the spatial interaction radius generates a larger number of competing transmission candidates and decreases the identifiability of individual infection pathways; this suggest that the results are more reliable using a larger cut-off distance. Notice that in most other papers there is no cut-off distance, but simply a declining transmission kernel; this probably explains why our estimate of *α* (the parameter controlling the decay of the kernel) is between 0.6 and 0.7, lower than the values estimated [14] or assumed [12] in similar models.

On the other hand, changing the value of *γ* (that controls the increase of infectiousness getting closer to the detection date) has only a minor effect on all results.

Overall, these findings indicate that the proposed framework successfully captures the main drivers of transmission while explicitly quantifying the uncertainty associated with epidemic reconstruction. Rather than identifying a unique transmission tree, the model provides a probabilistic ranking of plausible transmission pathways, allowing epidemiological interpretation to be combined with independent genetic evidence in order to assess the robustness of inferred transmission links.

### 4.2 Limitations and future directions

This study presents a mechanistic framework for reconstructing transmission pathways at the farm level; however, several limitations should be acknowledged. Infection times are never directly observed, and are instead inferred through the model. This introduces additional variability and may affect the identifiability of transmission pathways, particularly in cases with multiple plausible infectors.

First of all, although the infection probability matrix can be used to reconstruct the inferred transmission network, transmission probabilities are estimated conditional on the infection of each recipient farm rather than through a joint inference of the complete transmission tree. Consequently, the resulting network is not explicitly constrained to be globally consistent with the genetic relationships among all infected farms.

Indeed, the main focus of this study has been on the reconstruction of the most likely infectors for each farm to be compared with the genomic data when available. For this reason, we have studied the probabilities of the attribution of an infection to a specific transmission pathway rather than a statistically based estimate of parameter uncertainties.

Second, while our model uses the information on the company to which the farms belong, as this may be relevant for transmission, we have neglected other information, such as the size of the farms, and the type of birds raised. Although other studies have found that contact rates may depend significantly on these factors, adding more parameters to the model would lead to unidenfiability issues.

Similarly, data on contacts between farms via transport vehicles are available (including date, source, and destination, comprising approximately 35,000 recorded contacts) and could potentially provide additional information on transmission dynamics. In this work, we explored the possibility of incorporating these contacts into the transmission model as an additional network layer. However, introducing an explicit parameter regulating contact-based transmission led to identifiability issues and prevented stable convergence of the estimation procedure. Consequently, this component was not retained in the final model.

We also evaluated the consistency between recorded contacts and the inferred transmission pathways. In particular, observed contacts were compared both with the most likely inferred infector and with the set of the three most likely infectors having transmission probability greater than 0.1, while accounting for temporal compatibility between contact dates and estimated infection times. Overall, the agreement between the reconstructed transmission links and the available contact records was limited. For the most likely infector, only 4.3% and 3.4% of inferred transmission links corresponded to recorded contacts for *d*_max_ = 2 km and *d*_max_ = 1.5 km, respectively, with temporally plausible contacts accounting for 3.7% and 2.7% of cases. Considering instead the set of plausible infectors increased the agreement slightly: at least one recorded contact was identified for 7.4% of farms (*d*_max_ = 2 km) and 6.2% (*d*_max_ = 1.5 km), while temporally compatible contacts were found for 6.9% and 5.5% of farms, respectively. These results suggest that the available contact data alone are insufficient to explain the observed epidemic patterns. This discrepancy may reflect incomplete reporting, indirect transmission mechanisms not captured by direct movements, or temporal uncertainty in both the epidemiological reconstruction and the movement records.

Third, in our analysis we assumed a spatial cutoff (*d*_max_) for the distance-based infection force, and a temporal component for the force of infection from farms that increases (through the parameter *γ* getting closer to the detection time. The sensitivity analysis shows that the conclusions are very robust relatively to the value of *γ*, while the relative attribution of infections to the distance-based and company components depends to some degree on the value of *d*_max_, highlighting the importance of appropriately characterizing the effective spatial range of transmission.

Finally, the wildlife component is modeled as a homogeneous background infection pressure. This simplification does not capture possible spatial or temporal heterogeneity in wildlife-mediated transmission, which may become important in specific geographic areas or epidemic phases. Additional information on the movement of wildlife, either observed or estimated, might help narrowing down which farms are at risk at any given moment throughout the epidemic.

We remark that, while the comparison with genetic distances provided strong support for high-probability transmission links, the genetic information was not directly integrated into the inference procedure.

Future developments could therefore focus on incorporating genetic similarity directly within the likelihood framework, allowing epidemiological and molecular evidence to jointly contribute to transmission reconstruction. Integrating additional sources of information, including transport networks, pathogen genetic similarity, wildlife surveillance, within a unified Bayesian inference framework is an important goal for the future.

### 4.3 Implications for HPAI prevention and control

Although primarily developed for retrospective reconstruction, the proposed framework has relevant implications for HPAI prevention and control in densely populated poultry areas. The dominant contribution of spatial proximity is consistent with epidemiological and genomic analyses of the 2021–2022 Italian epidemic, which identified geographic distance, temporal compatibility between risk windows, and production-chain structure as relevant features of the observed spread [8, 30]. These findings suggest that, once HPAI is introduced into a poultry-dense area, subsequent epidemic spread may be driven predominantly by local farm-to-farm transmission. Reducing the duration and intensity of exposure to active outbreaks therefore requires timely detection, rapid depopulation and disposal, strict movement restrictions, and measures aimed at reducing local susceptibility during high-risk phases. The model could support retrospective assessment of the spatial scale most compatible with farm-to-farm transmission and exploration of alternative assumptions for restriction zones, enhanced surveillance areas, and pre-emptive measures. The importance of farm density, proximity, and biosecurity is also reflected in subsequent Italian legislation introducing minimum-distance requirements for the opening or reconversion of poultry farms and specific provisions for high-risk areas [31]. Although the present analysis does not evaluate these measures directly, it provides a basis for future assessments of density-related and spatially targeted control strategies.

The limited agreement between inferred transmission links and recorded transport contacts should not be interpreted as evidence that movements were unimportant. In [8], only a small fraction of retrieved movements met the temporal criteria for potentially productive contacts based on the overlap between the out-bound risk window of a possible source and the in-bound risk window of a potentially infected farm. This suggests that available contact-tracing data captured only part of the relevant transmission pathways. Incomplete or non-standardised reporting, indirect contacts, environmental contamination, and uncertainty in infection timing may all reduce correspondence between recorded movements and reconstructed links. Contact data should therefore be collected more completely, consistently, and promptly, ideally covering all inbound and outbound movements of vehicles, personnel, equipment, feed, and service providers. This should include visits that may not be perceived as high risk, such as those by maintenance workers, gas technicians, electricians, and plumbers. Harmonised digital records would facilitate their rapid use in epidemiological investigations and integration into analytical workflows during outbreak response.

The wildlife or unobserved-source component remains important for representing external infection pressure, particularly during the introduction phase or when no plausible farm-level source can be identified. However, its lower contribution relative to local spatial transmission suggests that repeated wildlife-mediated introductions alone were unlikely to explain epidemic expansion after the virus entered the poultry-dense area. Local spread may also have been amplified by operational constraints during periods of high incidence. During the 2021–2022 epidemic, saturation of culling and carcass-disposal capacity left multiple active outbreaks simultaneously present in close proximity, increasing opportunities for exposure and secondary transmission [8]. Early warning at the wild bird-domestic poultry interface therefore remains essential, but effective control of large epidemics also requires sufficient surge capacity for rapid depopulation and disposal, minimization of outbreak duration, and interruption of farm-to-farm transmission.

Finally, the comparison with genetic distances illustrates the value of combining mechanistic modelling with molecular evidence. The smaller genetic distances observed among high-probability inferred links provide independent support for the overall biological plausibility of the reconstructed pathways, although they do not confirm that individual links represent the true transmission events. The framework could therefore help prioritise epidemiological links for further investigation, support retrospective outbreak review, identify potential weaknesses in biosecurity or contact-tracing systems, and compare alternative control scenarios

## Acknowledgments

Mattia Sensi acknowledges the support of Fondazione Caritro (Cassa di Risparmio di Trento e Rovereto) through the Bando Post-Doc 2024 project “Modelli matematici di malattie infettive piu’ ospiti e popolazioni eterogenee: applicazioni all’influenza aviaria”.

## Data Availability Statement

The source code used to perform the analyses presented in this study is publicly available in the accompanying GitHub repository https://github.com/SaraSottile/AvianFluNetwork.git.

Owing to confidentiality restrictions, the original epidemiological, farm-level, and genetic datasets cannot be made publicly available. To ensure reproducibility of the computational workflow, the repository includes synthetic datasets that reproduce the structure and format of the original data and allow all scripts to be executed. The synthetic datasets preserve the data structure required by the analysis while containing no real farm identifiers, locations, or genetic information.

## A Sensitivity analysis

### A.1 Distribution of parameter estimates

Table 5 summarises the distribution of the estimated parameters across runs for the two different values of *d*_max_ and of *γ*.

**Table 5:** Summary statistics (mean, median, and standard deviation) of the parameter estimates across the 100 optimization runs.

| | Parameter | $d_{\max} = 2 \text{ km}$ | | | $d_{\max} = 1.5 \text{ km}$ | | |
| --- | --- | --- | --- | --- | --- | --- | --- |
|  |  | Median | Mean | SD | Median | Mean | SD |
| $\gamma = 0.1$ | $\beta_w$ | $1.41 \cdot 10^{-4}$ | $1.41 \cdot 10^{-4}$ | $1.40 \cdot 10^{-7}$ | $3.16 \cdot 10^{-4}$ | $3.16 \cdot 10^{-4}$ | $3.27 \cdot 10^{-7}$ |
| | $\beta_c$ | $7.12 \cdot 10^{-4}$ | $7.11 \cdot 10^{-4}$ | $5.36 \cdot 10^{-6}$ | $1.00 \cdot 10^{-3}$ | $9.96 \cdot 10^{-4}$ | $1.35 \cdot 10^{-5}$ |
| | $\beta_d$ | 110.64 | 111.61 | 8.86 | 140.68 | 149.35 | 25.32 |
| | $\alpha$ | 0.61 | 0.61 | $5.87 \cdot 10^{-2}$ | 0.62 | 0.57 | $1.28 \cdot 10^{-1}$ |
| | Length $E \rightarrow D$ | 6 | 7.83 | 5.83 | 7 | 7.94 | 6.02 |
| $\gamma = 0.2$ | $\beta_w$ | $1.19 \cdot 10^{-4}$ | $1.19 \cdot 10^{-4}$ | $1.74 \cdot 10^{-7}$ | $2.79 \cdot 10^{-4}$ | $2.79 \cdot 10^{-4}$ | $3.22 \cdot 10^{-6}$ |
| | $\beta_c$ | $6.64 \cdot 10^{-4}$ | $6.64 \cdot 10^{-4}$ | $3.48 \cdot 10^{-6}$ | $9.40 \cdot 10^{-4}$ | $9.47 \cdot 10^{-4}$ | $5.67 \cdot 10^{-5}$ |
| | $\beta_d$ | 87.87 | 87.89 | 4.77 | 108.29 | 104.38 | 21.36 |
| | $\alpha$ | 0.67 | 0.68 | $3.91 \cdot 10^{-2}$ | 0.71 | 2.55 | 12.74 |
| | Length $E \rightarrow D$ | 6 | 7.51 | 5.63 | 6 | 7.68 | 5.97 |

Decreasing *d*_max_ to 1.5 km results in an increased value of all transmission coefficients, and especially of that related to the wildlife component. On the other hand, increasing *γ* to 0.2 days^*−*1^ results in a decreased value of all transmission coefficients, less so that related to transmission in the same company network. Using *γ* = 0.2 instead of *γ* = 0.1 resulted in moderately higher estimates of the spatial decay parameter *α*; to better appreciate the difference, we plot in Figure 4 the value of the force of infection at a given distance from the source for the four combination of parameters (remember then that these values have to be multiplied by the expression dependent on the time before detection). As can be seen (in the right panel a log-log scale has been used to emphasise the differences) the estimates obtained are very similar for the different parameter combinations, showing the robustness of the estimates.

**Figure 4:**
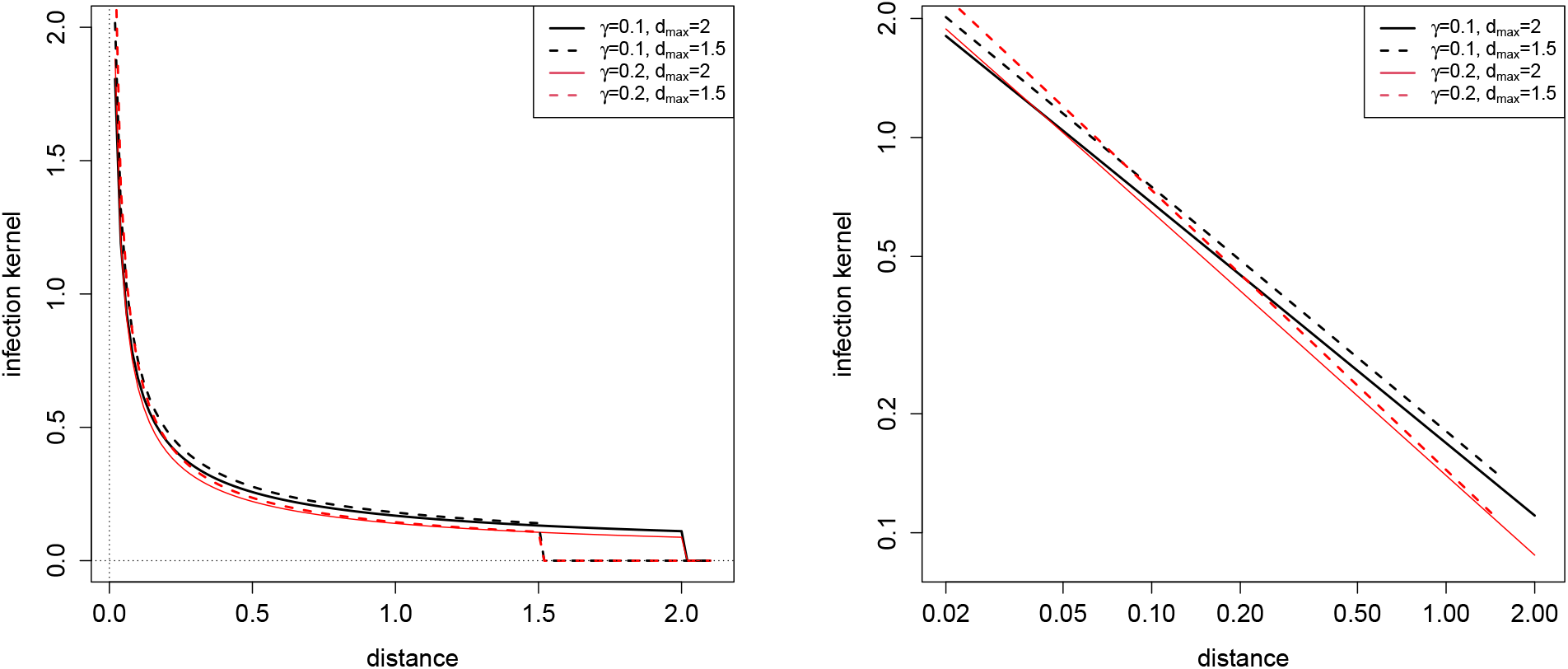
The estimated kernels showing how the force of infection depends on distance, for the four combinations of *d*_max_ and *γ* studied. Left: linear scale. Right: log-log scale.

The mean exposure-to-detection interval is slightly larger when *d*_max_ is decreased to 1.5 km, and is slightly smaller when *γ* is increased to 0.2 days^*−*1^, whereas the median is of 6 days for three of combinations, while increase to 7 6 days when *d*_max_ = 1.5 km and *γ* is 0.1 days^*−*1^. The differences in the mean and in the SD are modest, except for the configuration *γ* = 1*/*5 and *d*_max_ = 1.5 km. In that case, a small number (3%) of optimization runs converged to alternative local optima characterised by extreme values of *α*. We thus referred to the median only as a reliable indicator.

### A.2 Dominant transmission mechanisms

Table 6 reports the classification of the dominant transmission mechanism for the four combinations of *d*_max_ and *γ*. For each infected farm, the dominant mechanism is defined as the mechanism with the largest contribution to the estimated infection probability. The four categories distinguish infections attributed exclusively to farms belonging to the same company, exclusively to nearby farms, to farms that are both nearby and part of the same company, and to wildlife or other sources not explicitly represented in the dataset.

**Table 6:** Distribution of the dominant transmission mechanism for the four combinations of *γ* and *d*_max_. Values represent the number of infected farms assigned to each category.

| $d_{\max} = 1.5$ km | | | | |
| --- | --- | --- | --- | --- |
|  | Only same company | Only distance | Distance and same company | Wildlife |
| $\gamma = 0.1$ | 68 | 94 | 52 | 83 |
| $\gamma = 0.2$ | 77 | 91 | 51 | 78 |
| $d_{\max} = 2$ km | | | | |
|  | Only same company | Only distance | Distance and same company | Wildlife |
| $\gamma = 0.1$ | 59 | 124 | 64 | 50 |
| $\gamma = 0.2$ | 70 | 119 | 65 | 43 |

The results are broadly consistent across the two values of *γ*. For *γ* = 0.1, increasing the spatial cutoff from 1.5 to 2 km leads to an increase in the number of farms attributed to distance-based transmission, from 146 to 188, while the number attributed to wildlife decreases from 83 to 50. A similar pattern is observed for *γ* = 0.2, with distance-based transmission increasing from 142 to 184 farms and wildlife attribution decreasing from 78 to 43 farms. In contrast, the number of farms attributed exclusively to transmission within the same company changes less markedly, from 68 to 59 for *γ* = 0.1 and from 77 to 70 for *γ* = 0.2. The number of farms for which both distance and same-company transmission contributed to the dominant mechanism also remained relatively stable, changing from 52 to 64 for *γ* = 0.1 and from 51 to 65 for *γ* = 0.2.

The dominant mechanism was assigned with high probability in almost all cases. Only two farms had a dominant probability between 0.1 and 0.5: one for *γ* = 0.1 and *d*_max_ = 2 km, where both the distance-based network and the same-company network contributed to the dominant mechanism, and one for *γ* = 0.2 and *d*_max_ = 1.5 km, where the distance-based network alone contributed to the dominant mechanism. No farm had a dominant probability below 0.1.

Overall, the dominant mechanism is sensitive mainly to the assumed spatial cutoff, whereas the effect of *γ* is comparatively limited. A larger value of *d*_max_ shifts the attribution towards distance-based transmission and away from wildlife, while the relative contribution of same-company transmission remains comparatively stable. The high probability associated with the dominant mechanism in almost all cases also indicates that the classification of the dominant mechanism is generally well defined.

Table 7 reports the probability class of the most likely infector for the four combinations of *γ* and *d*_max_. Farm-to-farm infections were generally attributed to a specific infector with high probability, whereas the attribution to wildlife was more uncertain, with a larger proportion of cases falling in the Medium class. Low-probability assignments were observed only for *γ* = 0.2 and *d*_max_ = 2 km.

**Table 7:** Distribution of the probability class of the most likely infector according to network category, for the four combinations of *γ* and *d*_max_. Values represent the number of infected farms assigned to each probability class.

| $d_{\max}$ | $\gamma$ | Category | High | Medium | Low |
| --- | --- | --- | --- | --- | --- |
| 1.5 km | 0.1 | Only same company | 0 | 0 | 0 |
|  |  | Only distance | 92 | 3 | 0 |
|  |  | Distance and same company | 48 | 3 | 0 |
|  |  | Wildlife | 83 | 68 | 0 |
|  | 0.2 | Only same company | 0 | 0 | 0 |
|  |  | Only distance | 88 | 3 | 0 |
|  |  | Distance and same company | 50 | 1 | 0 |
|  |  | Wildlife | 78 | 77 | 0 |
| 2 km | 0.1 | Only same company | 0 | 0 | 0 |
|  |  | Only distance | 119 | 5 | 0 |
|  |  | Distance and same company | 61 | 3 | 0 |
|  |  | Wildlife | 50 | 59 | 0 |
|  | 0.2 | Only same company | 0 | 0 | 0 |
|  |  | Only distance | 117 | 4 | 0 |
|  |  | Distance and same company | 61 | 3 | 0 |
|  |  | Wildlife | 43 | 55 | 14 |

### A.3 Global comparison with genetic distances

The sensitivity analysis yielded results that are fully consistent with those obtained under the baseline parameters. As shown in Table 8, transmission pairs assigned to the high-probability class exhibited the smallest genetic distances for all parameter combinations, whereas medium- and low-probability pairs showed progressively larger genetic divergence. The same pattern was observed for the inferred most likely infectors, whose genetic distances closely matched those of the high-probability transmission pairs.

**Table 8:** Genetic distances stratified by inferred transmission probability for the two spatial cutoffs (*d*_max_ = 2 or 1.5 km) and *γ* = 0.1 or 0.2 (days)^*−*1^.

|  |  | High probability | Medium probability | Low probability | Most likely infector |
| --- | --- | --- | --- | --- | --- |
| $d_{\max} = 2, \gamma = 0.1$ | $n$ | 126 | 109 | 43,655 | 132 |
|  | Mean distance | 7.92 | 13.21 | 21.43 | 8.08 |
|  | Median distance | 5 | 11 | 21 | 5 |
| $d_{\max} = 1.5, \gamma = 0.1$ | $n$ | 101 | 80 | 43,709 | 104 |
|  | Mean distance | 7.84 | 13.15 | 21.41 | 7.90 |
|  | Median distance | 5 | 11 | 21 | 5 |
| $d_{\max} = 2, \gamma = 0.2$ | $n$ | 124 | 133 | 43,633 | 129 |
|  | Mean distance | 7.43 | 14.69 | 21.43 | 7.38 |
|  | Median distance | 5 | 14 | 21 | 5 |
| $d_{\max} = 1.5, \gamma = 0.2$ | $n$ | 97 | 119 | 43,674 | 99 |
|  | Mean distance | 7.54 | 16.03 | 21.41 | 7.46 |
|  | Median distance | 5 | 16 | 21 | 5 |

No significant global association between transmission probability and genetic distance was detected using Spearman’s rank correlation. Nevertheless, both the Mann–Whitney and the Kruskal–Wallis tests revealed highly significant differences among the probability classes for all parameter combinations (all *p <* 10^*−*35^), confirming that transmission pairs assigned higher probabilities were associated with significantly smaller genetic distances. The corresponding boxplots (Figure 5) clearly illustrate this trend. For all parameter combinations, the distributions shift progressively towards lower genetic distances from the low-to the high-probability class, while the genetic distances associated with the inferred most likely infectors almost completely overlap those of the high-probability transmission pairs. These findings further support the robustness of the inferred transmission network with respect to the assumed infectiousness profile.

**Figure 5:**
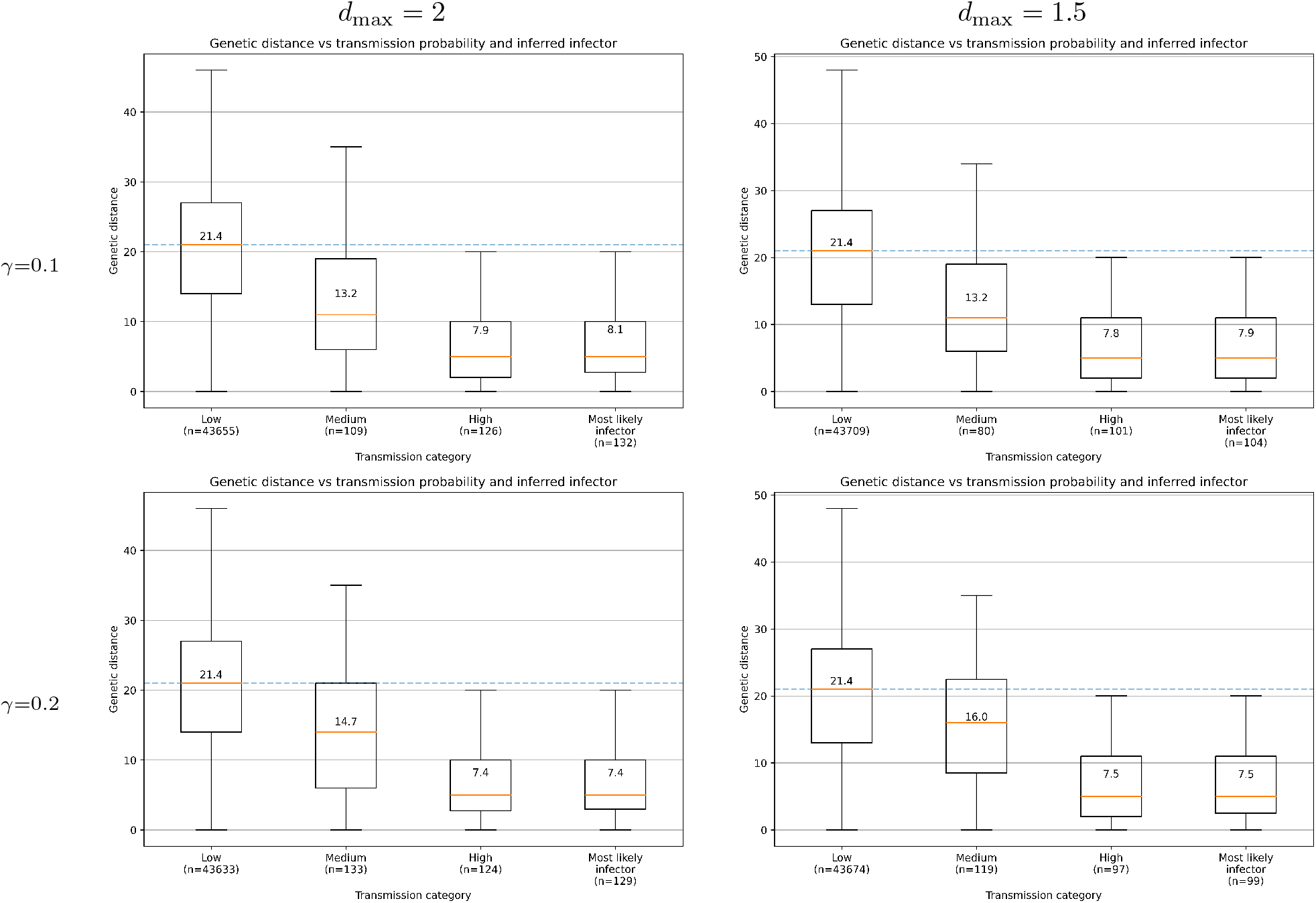
Distribution of pairwise genetic distances according to the inferred transmission probability classes under the sensitivity analysis.

## B Uncertainty in source attribution

As seen in the main text, the most likely infector has been identified for most farms with high probability (greater than 50%), but in some farms with medium probability (10 to 50%), and in a very limited number of farms (for some parameter combination only) the most likely infector had a probability below 10%.

Here, to quantify the uncertainty in attribution, we analysed the difference between the highest (*P*_1_) and second-highest (*P*_2_) transmission probabilities for each farm. The corresponding results are summarised in Table 9.

**Table 9:** Measures of uncertainty in source attribution for the two spatial cutoffs considered. Here, *P*_1_ and *P*_2_ denote the highest and second-highest transmission probabilities associated with each farm.

| | $\gamma = 0.1$ | | $\gamma = 0.2$ | |
| --- | --- | --- | --- | --- |
| | $d_{\max} = 2$ | $d_{\max} = 1.5$ | $d_{\max} = 2$ | $d_{\max} = 1.5$ |
| Mean gap ($P_1 - P_2$) | 0.510 | 0.440 | 0.533 | 0.453 |
| Median gap ($P_1 - P_2$) | 0.411 | 0.223 | 0.576 | 0.356 |
| $P_1 > 0.5$ | 60.6% | 47.1% | 59.9% | 46.5% |
| Gap $> 0.2$ | 60.9% | 55.9% | 61.3% | 55.6% |
| $P_1/P_2 > 2$ | 40.1% | 32.3% | 41.1% | 33.3% |

For all parameter combinations, a substantial fraction of farms exhibited a clearly dominant transmission source. When *d*_max_ = 2 km around 60% of farms have (for either value of *γ*) a most likely infector ascertained with probability exceeding 0.5, while the fraction of farms where the most likely infector has probability exceeding 0.5 decreases to around 47% with *d*_max_ = 1.5 km. A similar trend can be seen in all measures of uncertainties: mean and median gaps between the first and second most likely infectors or fraction of farms for which the ratio *P*_1_*/P*_2_ exceeds 2.

